# Single-camera, calibration-free gaze estimation using corneal reflections

**DOI:** 10.64898/2026.06.08.731021

**Authors:** David D. Au, Joshua B. Melander, Javier C. Weddington, Youssef Faragalla, Zaki Alaoui, Shenghua Liu, Qing Xu, Stephen A. Baccus

## Abstract

**Background:** Mice make substantial eye movements during head-fixed visual stimulation, and uncorrected gaze shifts corrupt receptive field measurements and confound stimulus-response relationships. Corneal-reflection video oculography in rodents has provided the methodological foundation for calibrated angular gaze tracking since Stahl (2004) but the calibration procedures used by existing methods — physical camera rotation, motorized stages, behavioral tasks, or precisely co-aligned dual cameras — have limited their adoption in many mouse neuroscience laboratories. Most studies instead use uncalibrated pupil tracking, deep learning pose estimation that returns pixel coordinates without angular calibration, or learned shifter networks that lack independent validation.

**New method:** We present an open-source corneal-reflection eye tracking system for head-fixed mice with two methodological contributions. First, a geometric model recovers gaze in calibrated angular units from the pixel displacements of the pupil and corneal reflections, using the known 3D positions of multiple fiducial LEDs as the source of angular scale. The model requires no estimate of *R*_*p*_, the per-animal eye-geometry parameter that earlier corneal-reflection methods determine through physical calibration. Second, a self-calibration procedure exploits the redundancy of multiple stationary fiducial LEDs: each LED produces an independent gaze estimate from the same geometric model, and a single residual calibration parameter is determined by minimizing the disagreement between per-LED estimates. This replaces the physical camera-rotation calibrations of earlier video oculography (Sakatani and Isa, 2004, 2007; van Alphen et al., 2013; Kretschmer et al., 2017), the motorized stages of Zoccolan et al. (2010), and the precision dual-camera alignment of Payne and Raymond (2017) with a software operation that requires no moving parts, no behavioral task, and no per-animal procedure. The system provides three interactive GUI stages: (1) pupil and LED detection via Difference-of-Gaussians filtering, (2) 3D geometry definition, and (3) gaze angle computation with blink detection, fiducial correction, and manual curation.

**Results:** Validation against a rotary-encoder-controlled artificial eye demonstrated mean absolute errors below 1° across all four fiducial LEDs over the ±20° working range of mouse eye movements, with Pearson correlations exceeding 0.998 between our method’s estimation and encoder ground truth. The self-calibration reduced inter-LED disagreement by a factor of 4–6 in mouse recordings. Gaze-corrected stimulus reconstruction applied to Neuropixels recordings from mouse V1 produced qualitatively sharper receptive field estimates with improved signal-to-noise ratios.

**Comparison with existing methods:** Our method is the first multi-LED, single-camera, fully software-calibrated corneal-reflection eye tracker for mice and includes an integrated open-source pipeline for detection, calibration, blink handling, and artifact correction. The multi-LED redundancy doubles as an internal consistency check — if two LEDs disagree on gaze direction, the calibration is wrong — providing a guarantee that learned approaches relying on neural-data-derived correction cannot offer.

**Conclusions:** Our method makes calibrated corneal-reflection eye tracking accessible to non-specialist mouse laboratories using consumer-grade hardware (∼ $2,000–2,700 USD), eliminates the per-animal calibration procedures of earlier methods, and is validated by two independent ground truths at both the absolute angular (artificial eye) and functional (V1 receptive fields) levels.

**Highlights:**

- Open-source corneal-reflection eye tracking for head-fixed mice using a single camera and multiple stationary fiducial LEDs.
- Geometric gaze model derives angular scale from LED positions, eliminating per-animal eye-geometry calibration.
- Self-calibration via multi-LED redundancy replaces physical camera rotation, motorized stages, and dual-camera precision alignment.
- Validated to sub-degree accuracy against a rotary-encoder ground truth across the ± 20° range of mouse eye movements.
- Gaze correction produces sharper V1 receptive field estimates in Neuropixels recordings.

## 1 Introduction

### 1.1 The need for calibrated gaze tracking in head-fixed mice

The head-fixed awake mouse has become a dominant preparation for studying visual processing, offering genetic tools, large-scale recording capabilities, and behavioral paradigms unavailable in primates. Despite this prevalence of the preparation, the implications of eye movements for mouse visual physiology have received comparatively little attention. Mice make substantial eye movements during visual stimulation, including saccades of up to ±20° in azimuth, slow drift, optokinetic responses, and positional shifts associated with locomotion and arousal state, which are increasingly recognized as behaviorally meaningful Sakatani and Isa (2004, 2007); Michaiel et al. (2020). These movements shift the retinal image and corrupt receptive field measurements, introduce variance into neural prediction models, and confound stimulus-response relationships.

Correcting for these shifts to estimate the retinal image requires knowing gaze direction in physical angular units (degrees), not pixel displacements. Despite this need, the majority of mouse neuroscience studies either ignore eye movements entirely or monitor only pupil position in un-calibrated pixel coordinates. This contrasts with primate work, where calibrated gaze estimates from the pupil-center / corneal reflection (P-CR) method enable precise receptive field mapping, stimulus-contingent experimental control, and post-hoc correction of retinal image motion as standard practice.

### 1.2 Existing approaches

Several methods exist for measuring eye movements in head-fixed mice, each with significant limitations for routine use. High-end systems — commercial trackers such as the Eyelink 1000 Plus (noa) and scleral search coil (Robinson, 1964) — provide sub-degree accuracy but are impractical at scale: commercial systems require behavioral calibration such as fixation that mice cannot perform and cost $25,000 or more, whereas search coils require surgical implantation and can impede natural eye movement. Deep-learning pose estimation frameworks like DeepLabCut (Mathis et al., 2018))and simpler pupil-thresholding tools (Syeda et al., 2024)) are accessible and widely deployed but output pixel coordinates rather than calibrated angles, and pixel-based tracking conflates eye rotation with translational artifacts. Learned shifter networks (Walker et al., 2019; Wang et al., 2025; Parker et al., 2022; Zahler et al., 2021) correct stimulus or world camera videos for eye movements using neural data, but the same neural data trains the shifter and validates it, an inherent circularity that mouse implementations cannot resolve without an independent gaze reference.

A line of work using corneal-reflection video oculography in rodents has provided the methodological foundation for calibrated angular gaze tracking in mice. Stahl (2004) introduced a single-camera, single-LED method in which gaze is recovered using *R*_*p*_, the distance from the pupil to the center of corneal curvature, as a per-animal calibration parameter determined by physically rotating the camera through known angles. Variants of this approach have been adopted in subsequent work with population-average *R*_*p*_ values (Sakatani and Isa, 2004, 2007) or refined calibration procedures (van Alphen et al., 2013; Kretschmer et al., 2017). Zoccolan et al. (2010) automated the calibration with motorized stages that move a single LED to known positions. Payne and Raymond (2017) bypass *R*_*p*_ estimation entirely using two precisely co-aligned cameras at a fixed angle, in service of validating a magnetic eye tracking system. These methods establish that calibrated angular gaze in rodents is achievable, but each requires procedures such as physical camera rotation, motorized hardware, or precision dual-camera alignment.

### 1.3 Contribution

This open-source corneal-reflection eye tracking method introduces two methodological contributions that together distinguish it from prior eye tracking in rodents. The first is a geometric model that recovers gaze in calibrated angular units from the pixel displacements of the pupil and corneal reflections, using only the known 3D positions of the fiducial LEDs as the source of angular scale. The model requires no estimate of *R*_*p*_, the per-animal parameter that underlies the Stahl (2004) calibration procedure and following work, and requires no information about the camera’s physical scale or intrinsic parameters.

The second contribution is a self-calibration procedure that exploits the redundancy of multiple stationary fiducial LEDs: each LED provides an independent gaze estimate from the same geometric model, and the small residual calibration parameter that the geometry alone cannot fix is determined by minimizing the disagreement between per-LED gaze estimates. This replaces the physical camera-rotation calibrations of (Stahl, 2004; Sakatani and Isa, 2004; van Alphen et al., 2013; Kretschmer et al., 2017) and motorized calibration stages of Zoccolan et al. (2010) with a software operation that requires no moving parts, no behavioral task, and no per-animal procedure. The multi-LED redundancy also serves as an internal consistency check for calibration that learned approaches relying on neural-data-derived correction cannot provide. Here we describe the system including the geometric model and self-calibration procedure, and validate the method with two independent measurements: absolute angular accuracy against a rotary-encoder-controlled artificial eye, and functional accuracy against improved receptive field estimates in mouse primary visual cortex.

## 2 Materials and Methods

### 2.1 Hardware

All components are commercially available, assembled without specialized mounting or motorized stages, and rigidly fixed for the duration of a recording session. The imaging system consists of an infrared-sensitive camera (Teledyne FLIR, Grasshopper3, 90 fps at max. resolution) focused on the mouse eye, two or more fiducial infrared LEDs positioned at known 3D locations around the stimulus display to produce corneal reflections, and infrared flood lamps that provide diffuse fill illumination for pupil contrast (Figure 1A). Fiducial LEDs and flood lamps operate at 850 nm to avoid interfering with visual stimulation. A zoom lens on the camera yields a field of view that captures the pupil and surrounding corneal reflections at high spatial resolution (Figure 1B). The LEDs are spaced with sufficient angular separation (∼ 30–70° as seen from the eye) to keep their corneal reflections spatially distinct across the full range of mouse eye movements and to provide the self-calibration procedure (Section 2.4) with multiple independent references for optimization. The complete hardware bill of materials, including part numbers, suppliers, approximate prices, and current computational requirements for the software, is provided in Supplementary Methods S1. Total hardware cost for the eye tracker components alone is approximately $2,000–2,700 USD.

**Figure 1:**
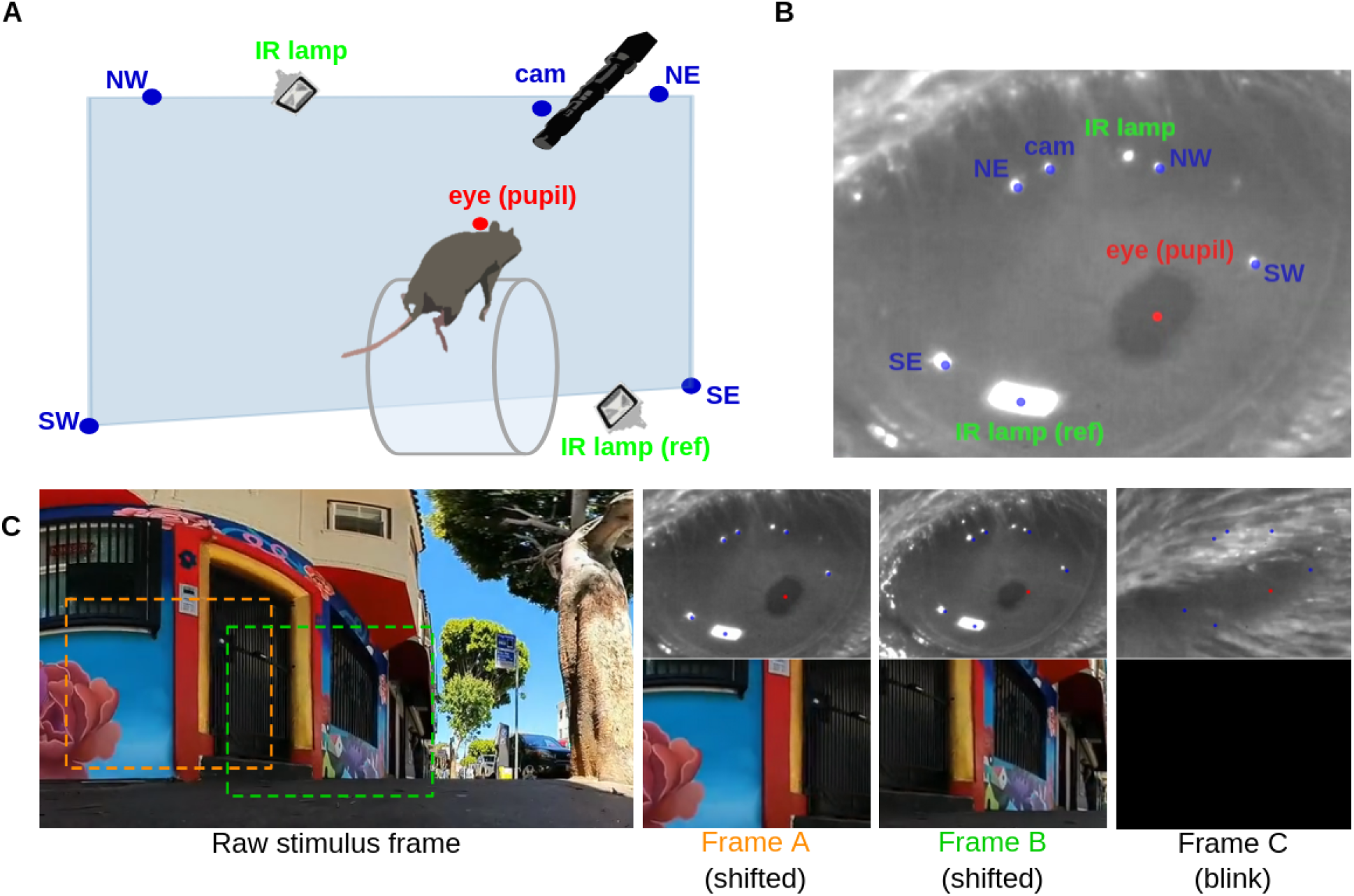
System overview and hardware. (A) Schematic of the head-fixed mouse recording setup showing the FLIR Grasshopper3 camera with Thorlabs zoom lens, 850 nm fiducial LEDs, and infrared flood lamps positioned around the mouse eye. (B) Example IR video frame with overlaid detections: pupil marker (red), fiducial and camera LED markers (blue), and floodlamp reflections (lower lamp reflection with blue marker is used for blink detection, higher lamp reflection is unmarked). (C) Raw stimulus frame with gaze-shifted and cropped example Frames A (orange dotted outline) and B (green dotted outline) that show different positional components after shifting, with Frame C representing a blacked-out blink frame.

#### 2.1.1 Stimulus display and synchronization

For visual electrophysiological experiments, visual stimuli were presented on a 24” 1920 *×* 1080 IPS monitor at 165 Hz, with a photodiode mounted on the monitor for temporal alignment to neural recordings. Analog signals from the photodiode, locomotion wheel, and camera strobe were acquired through a multi-channel USB DAQ. Neural activity was recorded using Neuropixels probes in mouse V1. Specifications and part numbers are listed in Supplementary Methods S1.

### 2.2 Coordinate systems

Geometric corneal-reflection eye tracking uses three coordinate systems (Figure 2A). The *world* system is a Cartesian system in which the user specifies the 3D positions of the eye, the camera, and each fiducial LED. The *image* system is the 2D pixel system of the camera sensor, in which the pupil and the corneal reflections of the LEDs and of the camera are located in each video frame. The *eye-centered* system is a spherical system centered on the eye, with its polar axis aligned along the line from eye to camera; angular position in this system is given by elevation and azimuth relative to that optical axis.

**Figure 2:**
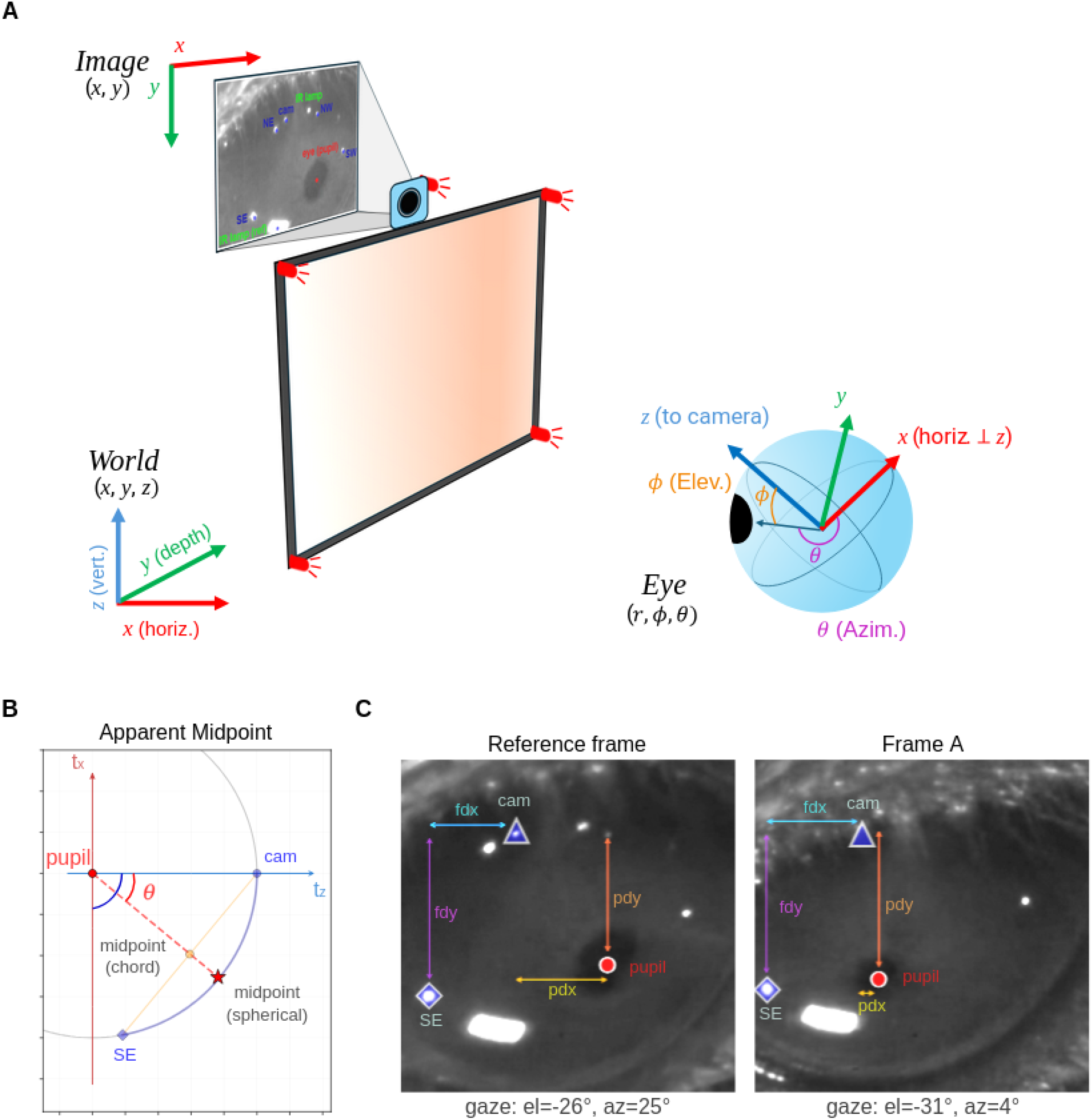
Gaze angle computation geometry. (A) Three-part coordinate system for gaze reconstruction between 3D World Cartesian (bottom left), 2D Image Cartesian pixel system (top left), and Eye-centered spherical system (right). (B) The apparent midpoint concept: the corneal reflection appears at the angular midpoint between the LED direction and the optical axis due to the convex mirror geometry of the cornea (focal length = *r/*2). (C) Two frames illustrate single fiducial LED pixel-to-angle mapping for the Reference frame and example Frame A with markings to indicate the pupil (red circle), SE LED reflection (blue diamond), and camera LED reflection (blue triangle). Displacement vectors *p*_*d*_ (pupil minus fiducial) and *f*_*d*_ (fiducial minus camera) are shown with their relationship to gaze angle approximations.

The gaze computation converts both input streams into the eye-centered system. The world positions of the camera and LEDs are recentered to the eye, rotated into the eye basis (Appendix A.2), and reparameterized as elevation and azimuth. These static eye-centered LED angles enter the geometric model of Section 2.3, which uses the per-frame image-system pixel positions of the pupil and corneal reflections to recover the eye-centered angular position of the pupil. That angular position is the gaze direction. When the result is needed in world coordinates — for example, to identify the point on the stimulus display being fixated — the eye-centered gaze direction is rotated back into the world basis by the inverse of the rotation used to convert the LEDs (Section 2.3.3, Appendix A.7), and the resulting 3D vector is then intersected with the display surface or re-parameterized as world-frame elevation and azimuth relative to the display normal.

### 2.3 Geometric model for gaze estimation

This section derives how the pixel positions of the pupil and corneal reflections are converted into the eye-centered angular position of the pupil. An IR LED at a fixed position produces a virtual image — the first Purkinje reflection — on the corneal surface. As the eye rotates, the pupil rotates in relation to the LED reflection, which moves only a small amount due to the change in corneal curvature. The displacement of the pupil relative to a corneal reflection is therefore a signature of eye rotation that is approximately invariant to translational motion of the eye in the image that can occur due to slight movement of the animal even in a head-fixed preparation. This property is the basis of all corneal-reflection eye tracking methods and motivates the pupil-CR vector as the gaze signal used in our method.

#### 2.3.1 The apparent midpoint

The corneal reflection of an LED does not appear at the LED’s angular position; it appears in the direction that bisects the angle between the LED and the camera as seen from the eye because the cornea acts as a convex mirror with focal length approximately equal to half the corneal radius of curvature. This places the virtual image of an LED at the midpoint of the chord between the points on the unit sphere in the LED direction and in the camera direction (Figure 2B).

The midpoint can be computed exactly. The LED direction and the optical axis are both unit vectors in the eye-centered Cartesian system; their chord midpoint is the average of the two vectors, and the angular position of that midpoint, recovered by reprojecting back to the unit sphere, gives the apparent angular position of the corneal reflection in the eye-centered system. For small angles, the apparent midpoint reduces to half the LED’s angular position, which is the approximation used in earlier corneal-reflection literature. The chord-midpoint computation is the exact form and is used throughout our method. The LED on the camera itself has its corneal reflection on the optical axis and serves as the zero-gaze reference. Additional details are given in Appendix A.5.

#### 2.3.2 Pixel-to-angle mapping

The pixel-to-angle mapping assumes that the camera sensor is oriented such that the image *y*-axis aligns with eye-centered elevation and the image *x*-axis with eye-centered azimuth. The image-system pixel positions of the pupil *P*, the corneal reflection *F* of a single LED, and the corneal reflection *C* of the camera enter the gaze computation as displacements. Both the pupil and the LED reflection are measured relative to the camera reflection, which represents the optical axis and serves as the zero-gaze reference:

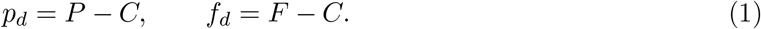

The pupil-to-camera displacement *p*_*d*_ encodes the gaze direction; the LED-reflection-to-camera displacement *f*_*d*_ encodes the apparent angular position of the LED’s corneal reflection in the image. Their ratio is dimensionless: when the pupil sits on the camera reflection, the ratio is zero and gaze is along the optical axis; when the pupil sits on the LED reflection, the ratio is one and gaze is in the direction of the LED.

Because the apparent midpoint of the LED in the eye-centered system is known (Section 2.3.1), the dimensionless ratio scales directly to an angular gaze position. For the elevation component:

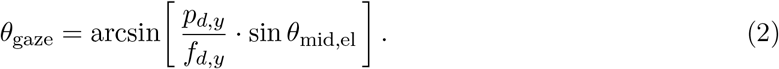

Then the azimuth component is computed with a cosine factor that corrects for horizontal foreshortening at non-zero elevation:

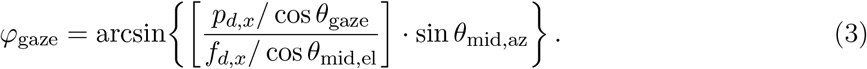

Both expressions are clipped to the valid range of arcsin to handle edge cases at extreme gaze angles. The full derivation is given in Appendix A.6.

This formulation has three properties worth noting. First, it returns gaze in calibrated angular units (degrees) without requiring the physical scale of the image — the pixel-to-degree mapping is set entirely by the eye-centered LED angles, which come from the user-supplied world geometry. Second, it does not require an estimate of *R*_*p*_, the distance from the pupil to the center of corneal curvature, which has been the central per-animal calibration parameter in earlier corneal-reflection methods (Stahl, 2004; Kretschmer et al., 2017; Sakatani and Isa, 2004). Third, the same pixel-to-angle mapping applies to every LED, since each LED’s apparent midpoint is known independently from the geometry.

#### 2.3.3 Gaze output in world coordinates

The gaze computation returns gaze direction in the eye-centered spherical coordinate system, as elevation and azimuth relative to the optical axis. This is the natural representation for many downstream analyses — for example, characterizing oculomotor behavior, comparing gaze across recording sessions with different camera placements, or quantifying eye movement statistics — because angular eye movements are described directly without reference to the external apparatus.

For analyses that require gaze in world coordinates — most importantly, identifying the point on the stimulus display at the center of gaze — we convert the eye-centered gaze direction back to the world system by inverting the rotation used to bring the LED positions into the eye-centered system. The inverse rotation is computed once per recording from the world-frame positions of the eye and the camera, and applied per video frame to the gaze direction. Both representations are written to the output file: eye-centered gaze is the primary output, and world-frame gaze is provided for downstream stimulus mapping. The full derivation of the inverse transform is given in Appendix A.7.

### 2.4 Self-calibration via multi-LED redundancy

The pixel-to-angle mapping in Section 2.3 takes the camera reflection *C* as the optical-axis reference. Although an LED attached to the camera produces this reflection, the LED is necessarily offset from the camera’s optical axis, so the detected reflection lies at a small pixel offset from the true optical-axis position. Gaze accuracy depends on the accuracy of that estimate.

With two or more fiducial LEDs visible in the same video frame, the pixel-to-angle mapping can be applied independently to each LED, producing one gaze estimate per LED. In the absence of detection noise and with perfectly known geometry, these per-LED estimates agree exactly. In practice, two sources of disagreement appear. The first is irreducible noise in the detected centroid positions, which limits the per-frame precision of any individual LED’s estimate and is reduced by averaging across LEDs. The second is the camera-reflection offset error: any error in the assumed camera-reflection position produces a coherent bias in all per-LED estimates whose magnitude depends on each LED’s angular position relative to the optical axis. This LED-dependence makes inter-LED disagreement an observable signature of the offset error. This section describes how our method uses that signature to estimate the camera reflection position from the recorded video.

#### 2.4.1 The calibration problem and prior approaches

Earlier corneal-reflection methods address the camera-reflection problem indirectly, by estimating an intermediate quantity called *R*_*p*_ — the distance from the pupil to the center of corneal curvature. With *R*_*p*_ known, gaze can be recovered from the pupil and a single LED reflection without needing the camera reflection at all. The Stahl-lineage approach (Stahl, 2004) estimates *R*_*p*_ per animal by physically rotating the camera through known angles and tracking the resulting motion of the pupil and CR. Subsequent methods adopted variants of this approach with assumed population-average values (Sakatani and Isa, 2004) or refined calibration procedures (Kretschmer et al., 2017). Zoccolan et al. (2010) automated the procedure with motorized stages that move a single LED to known positions. Payne and Raymond (2017) bypass *R*_*p*_ estimation entirely by combining pupil-CR displacements from two precisely co-aligned cameras at a fixed angle.

We take a different approach. Rather than calibrating *R*_*p*_, it treats the camera reflection position as the calibration target directly and exploits the redundancy of multiple stationary fiducial LEDs to determine it.

#### 2.4.2 Inter-LED disagreement as a calibration signal

With two or more fiducial LEDs visible in the same video frame, the pixel-to-angle mapping (Section 2.3.2) can be applied independently to each LED-camera pair, producing one gaze estimate per LED. If the camera reflection position is correct, the per-LED gaze estimates are consistent with one another (modulo small detection noise) because they describe the same underlying eye orientation. If the camera reflection position is wrong, the per-LED estimates disagree systematically: each LED’s gaze estimate is biased by an amount that depends on its angular position relative to the optical axis, so the disagreement between two LEDs is a function of the camera reflection error.

This dependence makes inter-LED disagreement an observable signature of the calibration error. With *N* LEDs visible, there are *N* (*N* −1)*/*2 pairs of per-LED gaze estimates, and the magnitude of the disagreement across these pairs quantifies the residual error. Minimizing this disagreement is equivalent to correcting the camera reflection position.

#### 2.4.3 Grid search procedure

The camera reflection position is parameterized as a 2D offset (Δ*x*, Δ*y*) in pixels from a reference position determined by the geometry. For each candidate offset, the pixel-to-angle mapping is applied to every LED, yielding *N* gaze estimates per frame. The mean absolute inter-LED disagreement is computed across all LED pairs and averaged over a calibration period (typically the full session, or a representative subset of frames). The offset that minimizes this disagreement is selected as the optimal calibration.

The optimization is performed sequentially: first the elevation component of the offset is determined by minimizing the inter-LED disagreement in elevation, then the azimuth component is determined by minimizing the disagreement in azimuth (Figure 3). This sequential structure exploits the fact that the elevation and azimuth components of the gaze computation are approximately decoupled under the alignment assumption stated in Section 2.3.2.

**Figure 3:**
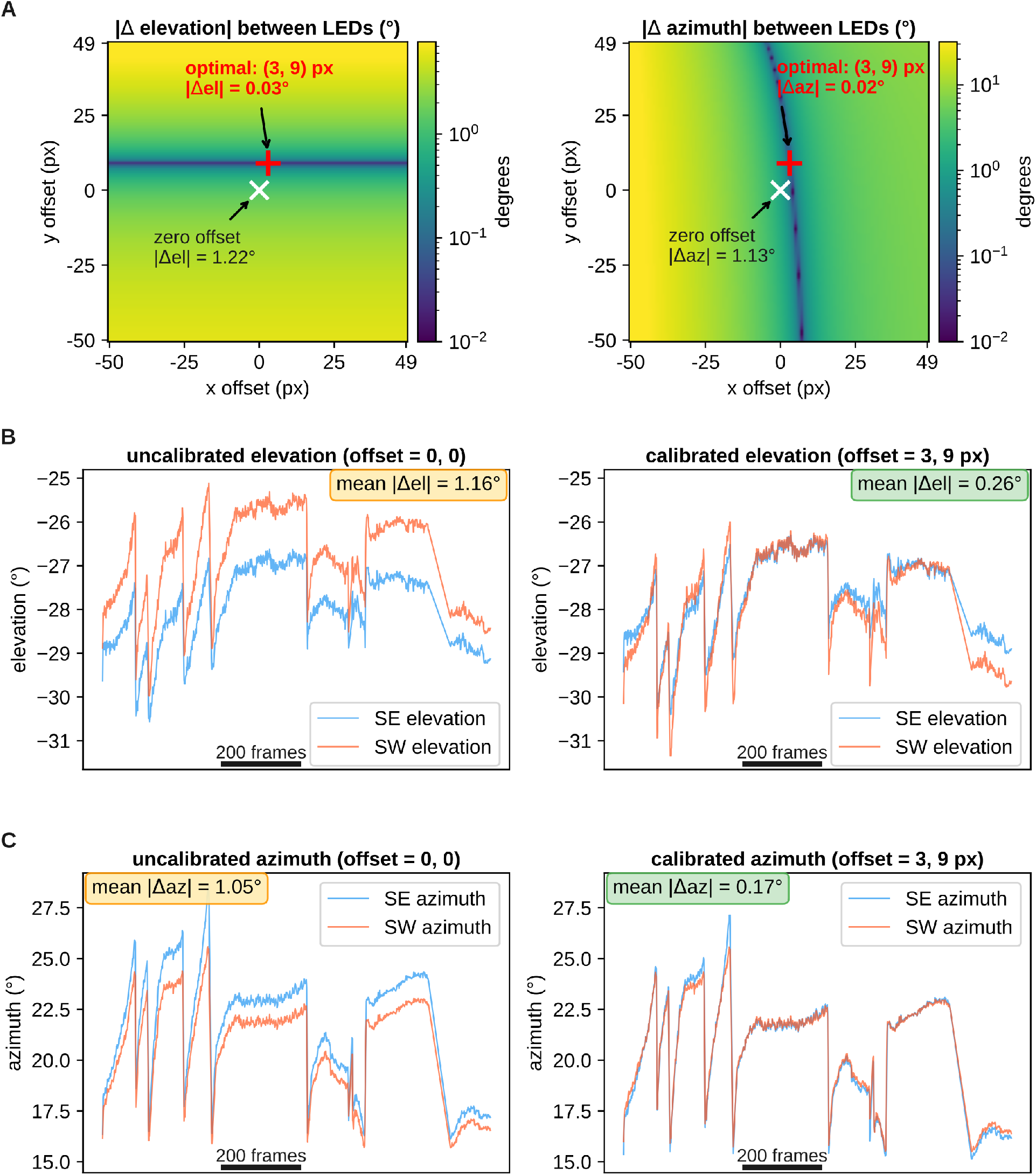
Grid search self-calibration. (A) Heatmaps of mean absolute inter-LED disagreement in elevation (left) and azimuth (right) as a function of camera offset (Δ*x*, Δ*y*). The clear minimum identifies the optimal calibration offset. Gaze traces from two LEDs (SE and SW) before calibration (showing systematic disagreement) and after calibration (showing convergence) for (B) elevation and (C) azimuth traces.

The grid is dense enough (sub-pixel resolution) to resolve the minimum to better than 1 pixel, and the search range is bounded by the plausible misalignment of the geometry-derived reference position. The full procedure operates on recorded images without hardware motion.

#### 2.4.4 Properties of the calibration

Several features of the calibration procedure are worth noting.

First, the calibration is *internal* : it uses only the recorded video in a session and the user-supplied geometry. No external ground truth, behavioral task, neural data, or auxiliary measurement is required. The inter-LED disagreement is computed from quantities that are already part of the gaze computation.

Second, the calibration is *self-validating* : if the optimal offset produces residual inter-LED disagreement that is large relative to the noise floor of centroid detection, the geometry is incorrect or the assumptions of the model are violated. Unlike learned gaze-correction approaches, which use neural data to train the correction, a failed calibration is therefore detectable from the data alone, without comparison to an independent measurement.

Third, the calibration is *overdetermined* when *N ≥* 3. With two LEDs the procedure is well-posed: one pair gives a 2D residual (elevation and azimuth disagreement) that constrains the 2D offset. With three or more LEDs the constraint is redundant, which improves robustness to detection noise on any single LED and allows direct estimation of calibration uncertainty from the spread of pairwise solutions.

### 2.5 Image processing and artifact handling

The pixel positions of the pupil and corneal reflections that enter the gaze computation are produced by an image-processing pipeline that runs once per video frame. The pipeline handles centroid detection, blink detection, and transient occlusion of individual fiducial reflections. None of these components are methodologically novel; they are described here at the level needed to interpret the gaze output, with implementation details in the appendix and supplementary material.

#### 2.5.1 Centroid detection

The pupil and each fiducial corneal reflection are detected as connected components in a difference-of-Gaussians (DoG) bandpass-filtered image. The DoG pipeline is applied separately to two channels: an inverted channel in which the dark pupil becomes a bright blob, and a direct channel in which bright LED reflections are detected. Connected components in the thresholded DoG output are filtered by area and by proximity to a user-defined reference point, and the centroid of the selected component is taken as the sub-pixel position. Default parameters and the full processing sequence are given in Supplementary Methods S3 and visualized in Supplementary Figure S1.

#### 2.5.2 Blink detection

Frames in which the eyelid occludes the cornea are detected by two complementary criteria, used independently or in combination depending on the recording configuration. The first criterion flags frames in which one or more fiducial reflections are displaced from their expected positions by more than a threshold, accompanied by a rapid change in detected pupil area. The second criterion uses the area of the brightest blob produced by the infrared flood lamp reflection on the eye surface: during blinks, the eyelid partially or fully occludes this reflection, reducing the blob area below a threshold. Detected blink intervals are excluded from the gaze computation. Both methods support post-hoc gap filling and boundary extrapolation; details are given in Supplementary Methods S3.

#### 2.5.3 Single-LED occlusion correction

When the eyelid or a transient reflection occludes one of the fiducial LEDs without producing a full blink, the affected LED’s gaze estimate is invalid for the duration of the occlusion. Our method handles this case by reconstructing the occluded LED’s pixel position from the visible LEDs, using an inter-LED pixel offset measured from frames in which all LEDs are simultaneously detected. The reconstructed position is treated as a measurement for purposes of the gaze computation in Section 2.3.2, with the caveat that the corresponding gaze estimate carries higher uncertainty. Frames in which two or more LEDs are simultaneously occluded are treated as blinks.

#### 2.5.4 Manual curation

The detection pipeline produces robust output across the large majority of frames in a typical recording, but edge cases — anomalous reflections from grooming, brief camera vibrations, partial lid closures that escape blink detection, and other rare events — are not fully handled by the automated pipeline. We implement a graphical interface for frame-by-frame inspection and editing of detected centroids and gaze trajectories, with tools for marking segment boundaries, labeling blinks, interpolating across gaps, and extrapolating at blink edges (Supplementary Methods S4). For the validation experiments in this paper, manual curation was applied to all sessions and accounted for a small fraction of total processing time.

## 3 Results

### 3.1 Sub-degree angular accuracy against rotary encoder ground truth

To establish the absolute angular accuracy independent of any neural or behavioral measurement, we constructed a physical surrogate eye with controllable orientation and measured the output against rotary-encoder ground truth. A polished metal sphere (∼ 8 mm diameter, approximating the mouse globe) was coated with titanium white acrylic paint to provide a diffuse reflective surface, with a small black circular pupil applied to the anterior face. The sphere was mounted on a precision rotary encoder and positioned at the location occupied by the mouse eye during normal recordings, preserving the camera–LED–eye geometry used in vivo (Figure 4A, B). A thin film of silicone oil was applied to the painted pupil and corneal-reflection regions to produce specular reflections detectable by the standard DoG pipeline.

**Figure 4:**
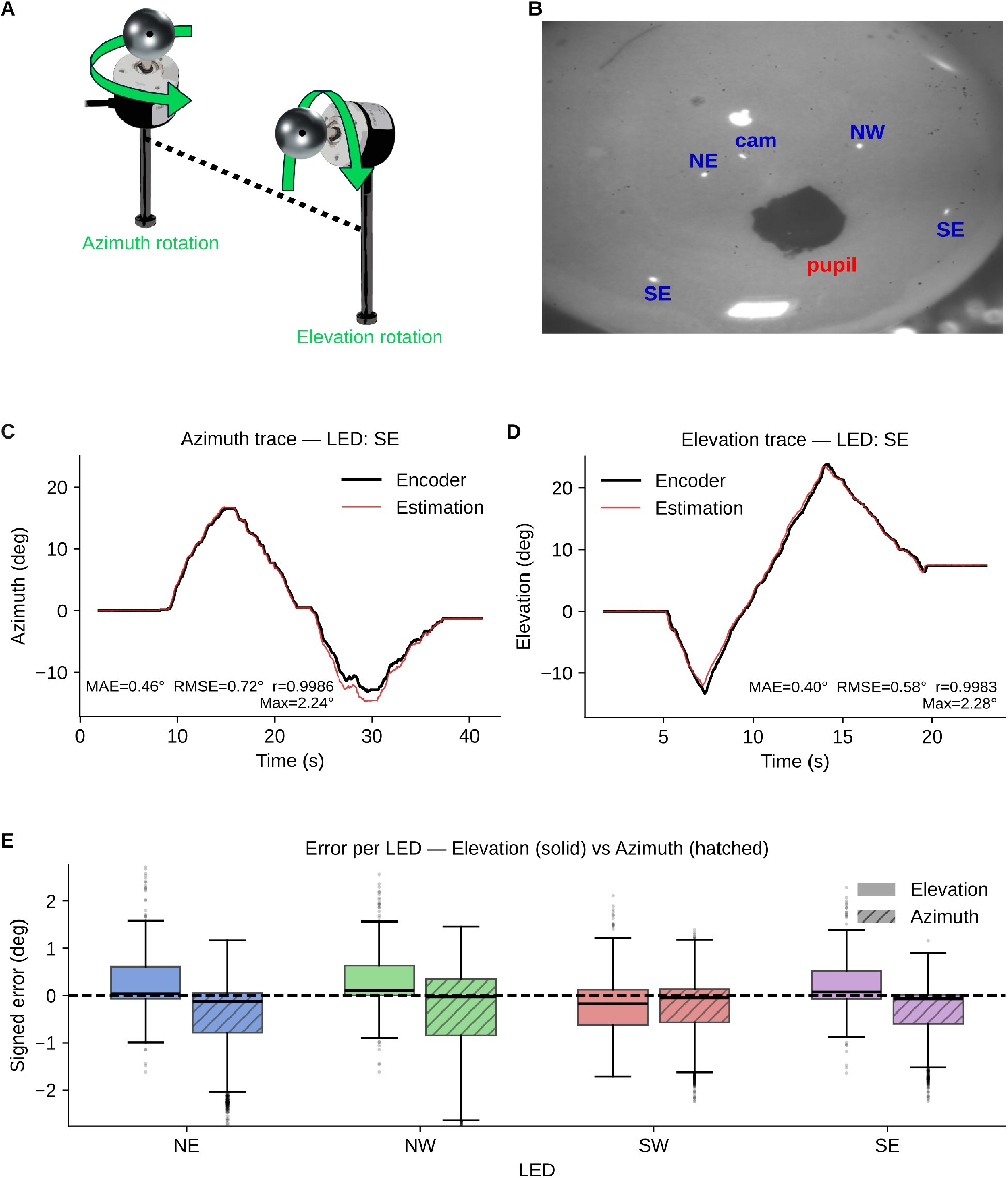
Artificial eye validation. (A) Schematic of the artificial eye setup where the head-fixed mouse is replaced with a metal sphere affixed to a rotary encoder that rotates in either azimuthal or elevational directions. (B) Example IR video frame from artificial eye with label annotations for the pupil and positional LED fiducials. (C) Azimuth and (D) Elevation traces comparing ground-truth rotary encoder angular displacement (black) overlaid with the geometric corneal-reflection estimation (red) from the SE LED. (E) Box plots of elevation (solid) and azimuth (hatched) error range boundaries with mean absolute error (MAE) for each positional LED.

The sphere was swept continuously through two ranges corresponding to the two gaze axes: − 10° to +24° in elevation, and −11° to +16° in azimuth. The encoder-reported angle served as ground truth, and our method’s gaze estimate was recorded independently from each of the four fiducial LEDs (NE, NW, SE, SW). We were able to track the encoder ground truth with high fidelity across both axes and all four LEDs (Figure 4C–E). The best-performing LED (SE) achieved mean absolute error (MAE) of 0.46°, root-mean-square error (RMSE) of 0.72°, and Pearson correlation *r* = 0.9986 for azimuth, with maximum absolute error of 2.24° (Figure 4C). Elevation estimation from the same LED was comparably accurate: MAE = 0.40°, RMSE = 0.58°, *r* = 0.9983, maximum error 2.28° (Figure 4D). Reconstructed gaze traces were nearly indistinguishable from the encoder trace across the full angular range, including smooth continuous sweeps and direction reversals.

Sub-degree accuracy was observed across all four LEDs, not only the best-performing one (Figure 4E). For azimuth, MAE was below ± 1° for every LED, with most error mass concentrated within 1° and tails extending to approximately ± 3° only at the extremes of the encoder sweep. Elevation errors showed similarly compact distributions, with all LEDs achieving MAE below ± 1° and error spread predominantly within ± 1°. The SE LED showed the smallest bias and tightest error distribution across both axes in this geometry, consistent with its favorable angular position relative to the camera axis; no systematic directional bias was apparent across the other LEDs.

These results establish a hardware- and model-level accuracy bound for geometric corneal-reflection eye-tracking that is fully independent of the neural data used in the functional validation. The sub-degree MAE achieved across all four LEDs confirms that the geometric model and grid-search self-calibration recover absolute gaze angles from corneal-reflection geometry at the level required for mouse V1 receptive field work.

### 3.2 Self-calibration convergence

The artificial-eye validation provides absolute angular accuracy with no reliance on neural data, but it does not directly test the self-calibration component of the pipeline, since the camera-reflection offset for the artificial-eye rig was determined by the same procedure used in vivo. To characterize the calibration itself, we examined the convergence of the grid search and the residual inter-LED disagreement that remains after calibration in mouse recordings.

The grid search over camera-reflection offsets (Δ*x*, Δ*y*) produced clear minima in inter-LED disagreement for both elevation and azimuth, well separated from the search-space boundaries (Figure 3A). The optimal offset was reproducibly identified to sub-pixel resolution. Before calibration, gaze traces computed independently from the SE and SW LEDs showed systematic offsets of ∼1° in elevation and ∼1° in azimuth (mean |Δel| = 1.16°, mean |Δaz| = 1.05°; Figure 3B, C, left). After calibration, the same gaze traces converged to within ∼0.2–0.3° of each other (mean |Δel| = 0.26°, mean |Δaz| = 0.17°; Figure 3B, C, right) — a reduction of ∼4–6*×* in inter-LED disagreement.

The residual inter-LED disagreement after calibration is comparable to the per-frame detection noise of individual centroids (corresponding to ∼0.1–0.3° at typical pixel resolutions), indicating that the calibration has reduced the systematic camera-reflection offset error below the noise floor of the rest of the pipeline. This residual is a quantitative diagnostic: large residuals after calibration would indicate that the geometry is incorrect or that the model’s assumptions are violated, and the calibration procedure therefore serves as a continuous self-check of the gaze output without requiring an independent measurement.

### 3.3 Improved V1 receptive field estimates with gaze correction

To validate that our method’s gaze estimates capture eye movements at sufficient accuracy to improve real neural data, we applied gaze-corrected stimulus reconstruction to Neuropixels recordings from mouse primary visual cortex during white-noise visual stimulation. For each video frame, the per-frame gaze direction was converted to a pixel shift in the stimulus coordinate system. A cropped window corresponding to each neuron’s receptive field region was extracted from each stimulus frame at either a fixed position (“static” reconstruction, ignoring eye movements) or a gaze-shifted position (“shifted” reconstruction, correcting frame-by-frame for the eye position). Blink frames were excluded from both reconstructions. Spike-triggered averages (STAs) were computed for each V1 neuron using both reconstruction methods.

Gaze-corrected stimulus reconstruction produced sharper and more spatially compact receptive field estimates compared to uncorrected reconstruction (Figure 5). Two representative neurons illustrate this effect: both exhibited clearly defined OFF subregions under gaze-corrected reconstruction, with peak STA *z*-scores of 10.57 and 10.87 respectively, compared to 5.85 and 4.03 under the uncorrected condition — a 1.8- to 2.7-fold improvement in signal-to-noise ratio. The uncorrected STAs exhibited spatially diffuse response regions consistent with the smearing predicted by uncompensated retinal-image motion, as quantified by the gaze position distribution (Figure 6). These results confirm that gaze estimates, validated above against an absolute physical ground truth, also operate above the accuracy threshold needed for meaningful visual receptive field measurements in mouse V1.

**Figure 5:**
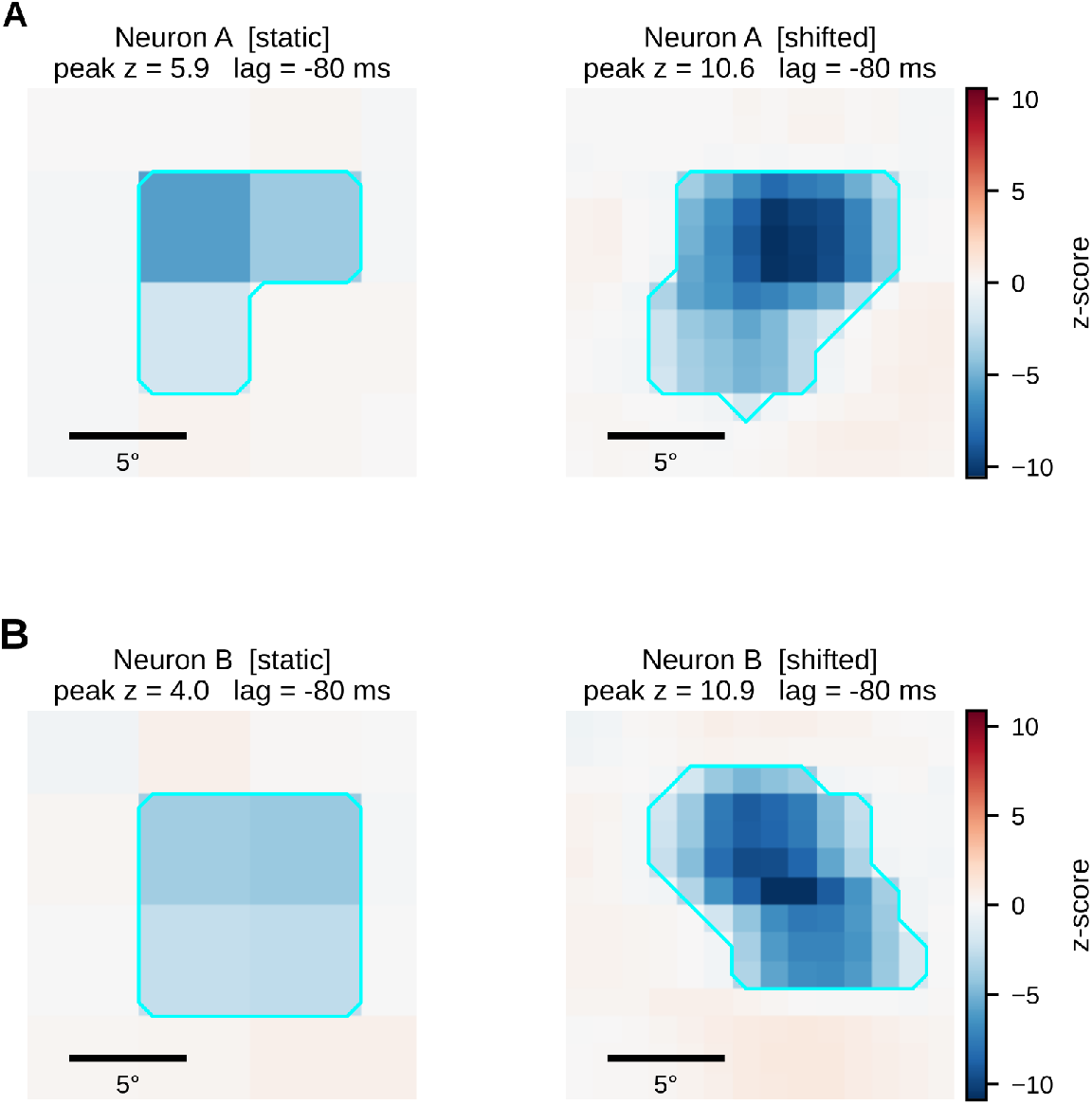
Gaze correction sharpens receptive field estimates from spike-triggered averaging. Peak frames of the spike-triggered average (STA) for two example neurons (A, B) computed from white noise stimuli under uncorrected (left) and gaze-corrected (right) conditions. In the uncorrected condition, the stimulus is treated as fixed on the screen regardless of the animal’s gaze. In the gaze-corrected condition, each stimulus frame is shifted to account for the instantaneous gaze position, stabilizing the stimulus in retinal coordinates. Cyan contours indicate the OFF receptive field region. Scale bar: 5°. Peak STA *z*-score is indicated for each panel.

**Figure 6:**
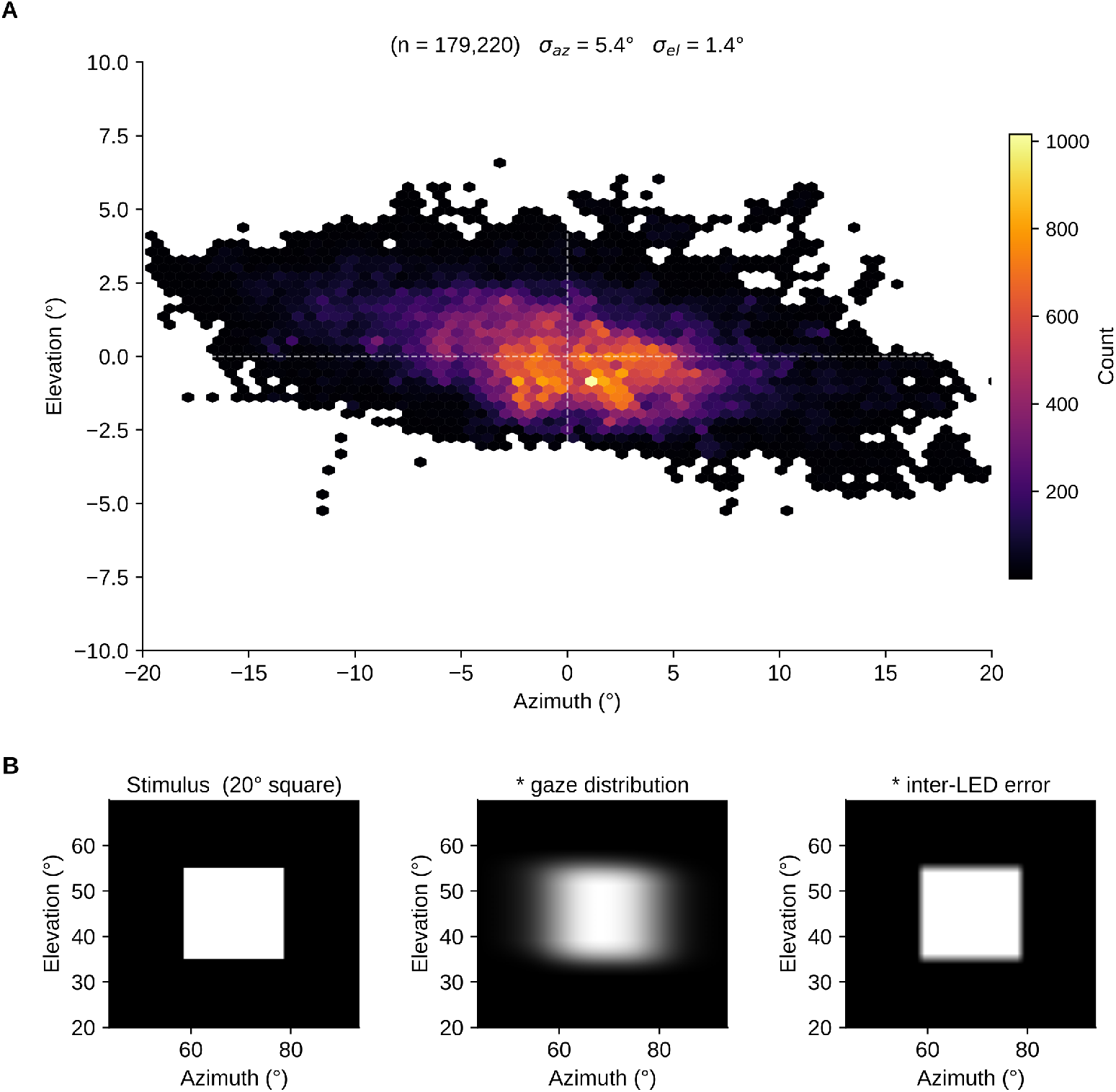
Eye position distribution and effective stimulus from an example dataset. (A) Hexbin density of gaze position (azimuth and elevation, mean-subtracted) across all valid frames (*n* = 179,220; *σ*_az_ = 5.4°, *σ*_el_ = 1.4°). (B) The 20° square stimulus presented at screen center. (C) Stimulus convolved with the gaze position distribution, showing the effective retinal stimulus location given the animal’s eye movements. (D) Stimulus convolved with the inter-LED error distribution, showing the spatial blur introduced by gaze estimation error. All panels share the same angular scale.

### 3.4 Gaze-correction recovers stimulus alignment obscured by natural eye movements

Analysis of gaze trajectories from a head-fixed recording session (*n* = 179,220 valid frames) revealed substantial natural eye movements during stimulus presentation (Figure 6A). Eye position was broadly distributed along the azimuth axis (*σ*_az_ = 5.4°, range − 20.5° to +25.6°) and more constrained in elevation (*σ*_el_ = 1.4°, range − 5.5° to +19.8°), consistent with the predominantly horizontal scanning behaviour observed in head-fixed mice. Convolving the nominal stimulus (20° square; Figure 6B, left) with the gaze position distribution illustrates the effective retinal stimulus in the absence of gaze correction: the sharp stimulus boundary is substantially smeared, particularly along the azimuth axis, producing a spatially diffuse input to the retina (Figure 6B, middle). By contrast, convolution with the inter-LED estimation error distribution, which reflects the residual uncertainty in our method’s gaze estimates (*σ*_az_ = 0.25°, *σ*_el_ = 0.42°), produces a negligible blurring of the stimulus (Figure 6B, right). The estimation error was 21.0 *×*and 3.2*×* smaller than the gaze variation in azimuth and elevation, respectively.

## 4 Discussion

### 4.1 Principal findings

We have presented geometric corneal-reflection eye-tracking as an open-source system for head-fixed animals with two primary methodological contributions. The geometric model recovers gaze in calibrated angular units from the pixel displacements of the pupil and corneal reflections, using the known 3D positions of multiple fiducial LEDs as the source of angular scale; it requires no estimate of *R*_*p*_, the per-animal eye geometry parameter underlying previous calibrations (Stahl, 2004), and no information about the camera’s physical scale. In addition, the self-calibration procedure exploits the redundancy of multiple stationary fiducial LEDs to determine the residual calibration parameter via a software grid search over inter-LED disagreement, replacing the physical camera rotations, motorized stages, and dual-camera alignments used by earlier corneal-reflection methods. Validation against a rotary-encoder-controlled artificial eye demonstrated mean absolute errors below 1° across the ± 20° working range of mouse eye movements, and gaze-corrected stimulus reconstruction produced sharper receptive field estimates in mouse V1 across a population of recorded neurons.

### 4.2 Comparison with prior methods

Our method belongs to a line of corneal-reflection video oculography methods in rodents beginning with Stahl (2004). The defining methodological choice across this lineage has been how to determine the parameters that convert measured pixel displacements into angular gaze. Earlier methods use *R*_*p*_ — the distance from the pupil to the center of corneal curvature — as the calibration parameter and determine it via physical procedures: Stahl (2004), Sakatani and Isa (2004), van Alphen et al. (2013), and Kretschmer et al. (2017) all rotate the camera through known angles to recover *R*_*p*_, while Zoccolan et al. (2010) automate the procedure using motorized stages that translate a single LED to known positions. These approaches share two practical constraints: they require per-animal or per-rig procedures that scale poorly to large cohorts, and they require precise physical control of either the camera or the calibration source, which limits adoption in laboratories without optical engineering expertise. Payne and Raymond (2017) bypass *R*_*p*_ estimation entirely with a dual-camera trigonometric configuration that determines angular scale from the known angle between two precisely co-aligned cameras; this is the closest precedent to geometry-based eye tracking approaches in eliminating *R*_*p*_ from the calibration. Our method achieves the same *R*_*p*_-free recovery with a single camera and multiple fiducial LEDs at known 3D positions, replacing dual-camera precision alignment with multi-LED redundancy and a software calibration procedure.

Geometric corneal-reflection eye-tracking also differs from non-CR approaches that have gained traction in mouse work. Deep-learning pose estimation (Mathis et al. (2018) and simple pupillometry tools can track the pupil with high pixel precision but do not provide angular calibration; converting their output to gaze in degrees requires additional steps that are rarely performed rigorously. Learned shifter networks (Parker et al., 2022; Zahler et al., 2021; Walker et al., 2019; Wang et al., 2025) correct stimulus or world-camera videos using neural data, but the same neural data trains and validates the correction, thus lacking independent validation. Geometric calibration avoids this circularity by deriving gaze from optical physics rather than from the data the gaze estimates are intended to correct. The multi-LED redundancy provides a direct internal consistency check: large residual inter-LED disagreement after calibration indicates that the geometry or the optical model is wrong, and the system therefore detects calibration failures from the recorded video alone without requiring an independent reference.

For applications targeting higher precision than our method provides — fixational eye movements, microsaccade dynamics, smooth-pursuit kinematics — dual Purkinje image methods such as OpenIrisDPI ((Ressmeyer et al., 2025; Wu et al., 2023)) achieve sub-arcminute precision in primates at the cost of a narrow tracking range (∼±10°). Geometric corneal-reflection eye-tracking and OpenIrisDPI are complementary: OpenIrisDPI for high-precision primate fixational studies, and our method for routine mouse gaze correction in visual physiology over a greater than ±20° range. This geometric model and self-calibration approach are species-agnostic and could in principle be applied to primates without modification, particularly for preparations in which behavioral fixation training is not feasible — marmosets, untrained adults, or high-throughput screening protocols — though commercial trackers and dual Purkinje image systems remain the appropriate choice for trained head-fixed macaque precision work.

### 4.3 Limitations and sources of error

Several limitations of the current method should be noted and group naturally into operational constraints and quantified error sources.

#### Operational constraints

Our implementation of geometric corneal-reflection eye-tracking assumes rigid head fixation: any movement of the camera or LEDs relative to the head-fixed eye during a session invalidates the geometric calibration. The system tracks two-dimensional gaze (elevation and azimuth) but not torsional eye rotation, which would require additional features on the eye such as the fluorescent markers used by van Alphen et al. (2013). At extreme gaze angles where the pupil is fully occluded by the eyelid or one or more fiducial reflections leave the cornea, gaze cannot be recovered; this becomes a practical limit beyond approximately ±25° in our rig geometry. The manual curation step required to handle anomalous frames (grooming reflections, brief camera vibrations, partial lid closures) constrains throughput for very large datasets, though it accounts for only a small fraction of total processing time in typical sessions.

#### Quantified error sources

The paraxial approximation in the apparent-midpoint model is exact in the chord-midpoint computation but introduces a small gaze-angle-dependent systematic bias when combined with the pixel-to-angle mapping under the alignment assumption stated in Section 2.3.2. This bias is below 1° within the central ± 20° range and grows to roughly 2–9° at 40° gaze, with magnitude and sign depending on which LED is used. For the majority of mouse visual physiology applications, where eye movements rarely exceed ± 20°, this bias is below the accuracy threshold needed for receptive field correction. The user-supplied geometry contributes a systematic angular uncertainty that scales inversely with LED-to-eye distance: at the ± 1 mm measurement tolerance and typical 10–25 cm LED distances, this corresponds to ∼ 0.2–0.6° of angular uncertainty in the LED positions, well below the per-frame detection noise of centroid localization. Detection noise in centroid positions sets the ultimate per-frame precision floor at ∼ 0.1–0.3° in our recording configuration.

#### Minimum operational luminance

While infrared illumination is unaffected by monitor state and provides stable corneal reflections across all conditions, the degree of pupil dilation at low visible luminance posed a challenge for the tracking algorithm. At larger pupil diameters, such as those observed during prolonged dim gray and black screen periods, the reduced contrast between the pupil boundary and the surrounding iris caused the tracking algorithm to fail more frequently. As a result, reliable eye tracking required a minimum level of visible light from the monitor to maintain sufficient pupil constriction. This represents a limitation of the current approach: gaze tracking data are most reliable during active stimulus presentation (14.2–50.4 µW/cm^2^) and may be degraded or unavailable during dark (0.25 µW/cm^2^) or low-luminance intervals such as inter-stimulus periods (0.59 µW/cm^2^).

#### Camera leveling requirement

The current implementation requires the camera to be physically leveled parallel to the table surface prior to recording, such that the camera’s horizontal and vertical image axes are aligned with the lateral and vertical axes of the world coordinate system. If the camera is rolled about its optical axis — even by a small angle — azimuth and elevation estimates become mutually contaminated: a gaze displacement that is purely horizontal in the world frame projects onto both pixel axes in the camera image, introducing crosstalk between the two angular components. This roll-induced coupling is not corrected for in software. In practice, leveling is achieved manually using a bubble level, which introduces some residual angular error. Future implementations could incorporate an explicit roll correction by estimating the camera’s in-plane rotation from the known world-frame geometry of the LED fiducials.

### 4.4 Conclusions and outlook

Applying geometry with corneal-reflections provides calibrated angular eye tracking for head-fixed mice using a single camera, multiple stationary fiducial LEDs, and a software self-calibration procedure that requires no per-animal physical calibration, no behavioral fixation task, and no motorized hardware. The two methodological contributions — the geometric model that derives angular scale from multi-LED geometry without *R*_*p*_, and the multi-LED self-calibration that replaces physical calibration procedures — together make calibrated corneal-reflection eye tracking accessible to mouse laboratories that have been constrained by the per-animal procedures of earlier methods. The validation against an artificial-eye rotary-encoder ground truth and the demonstration of improved V1 receptive field estimates establish the method’s accuracy at both the absolute angular and functional levels on independent data.

Several extensions are natural directions for future work. Deep-learning centroid detection (e.g., YOLO-based segmentation) could reduce the manual curation requirement while leaving the geometric calibration framework unchanged. Real-time gaze estimation would enable gaze-contingent stimulus paradigms, which are standard in primate work but rare in mouse studies. Automated geometry measurement using structured light or stereo vision could eliminate the manual ± 1 mm measurement step. Calibrated pupil diameter extraction from the existing blob detection pipeline would provide simultaneous gaze and arousal/attention measurements. Extending the geometric model to recover torsional eye rotation, using additional features such as iris landmarks rather than applied markers, would broaden the method’s applicability to oculomotor research. By releasing the software and hardware specifications publicly, we intend to lower the barrier to calibrated mouse eye tracking and to facilitate community-driven extensions.

## Acknowledgments

We would like to thank Robert Schneeveis for invaluable assistance designing, machining, and operating custom experimental equipment used in this manuscript.

## Data and code availability

An implementation of the methods described here is freely available at https://github.com/baccuslab/btrack/. The repository includes installation instructions, example data, geometry files, and Jupyter notebook tutorials. Raw data supporting the electrophysiological validation will be deposited upon publication.

## A Appendix Coordinate system conventions and derivations

This appendix provides the full mathematical derivation of our method’s coordinate transformations, illustrated with a worked example using a real experimental dataset and geometry file.

### A.1 World coordinate system

All positions are measured in centimeters in a right-handed world coordinate system: *x* = lateral (left/right across the table), *y* = depth (toward/away from the display), *z* = vertical (up from the table surface). The eye position *E* defines the origin after centering. For the geometry of the authors’ main experimental apparatus:

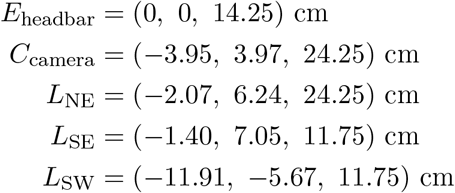

After centering (subtracting E):

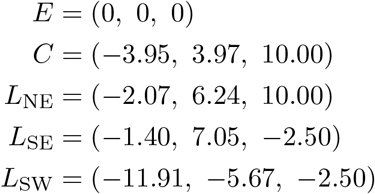

### A.2 Eye basis construction

The eye basis 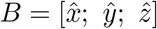 is constructed from the centered camera position *C*.

#### Step 1

The *z*-axis points from eye to camera:

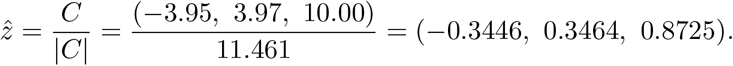

#### Step 2

Project *z* onto the horizontal plane (zero the vertical component):

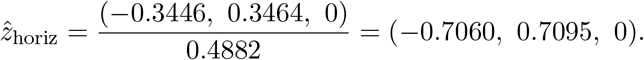

#### Step 3

The *x*-axis is horizontal and perpendicular to *z*:

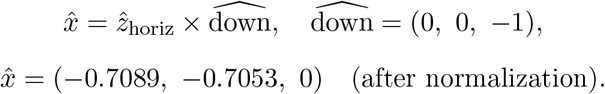

#### Step 4

The *y*-axis completes the right-handed system:

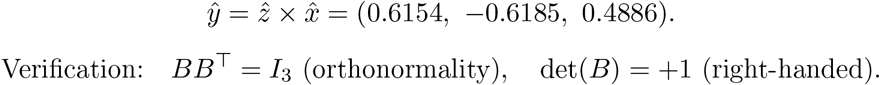

Verification: *BB*^*T*^ = *I*_3_ (orthonormality), det(*B*) = +1 (right-handed).

### A.3 World-to-eye coordinate transform

A point *P* in world coordinates is transformed to eye-centered Cartesian coordinates as *P*_eye_ = *P*_world_ *B*^*T*^. Since *B* is orthonormal, *B*^−1^ = *B*^*T*^. The components (*t*_*x*_, *t*_*y*_, *t*_*z*_) represent the position along the eye’s *x, y*, and *z* axes respectively. Example: the North East (NE) LED in eye coordinates:

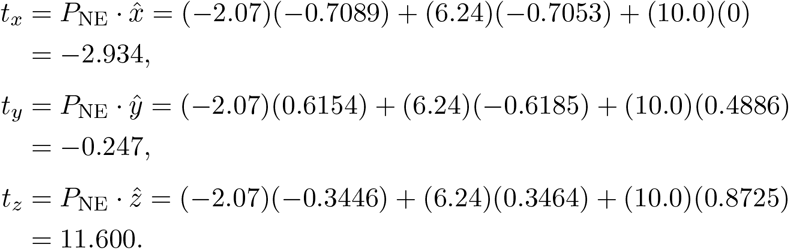

### A.4 Eye-centered spherical coordinates

The LED direction in eye-centered spherical coordinates (elevation, azimuth) is:

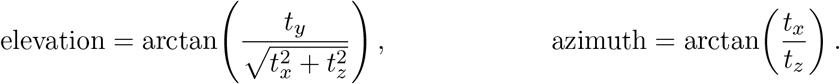

For the NE LED:

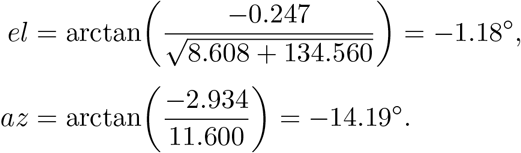

*Convention:* positive elevation is upward (toward +*y* in the eye frame), positive azimuth is leftward (toward +*x* in the eye frame).

### A.5 Apparent midpoint

The corneal reflection of an LED appears at the angular midpoint between the LED direction and the camera axis (the *z*-axis of the eye frame). Because the cornea acts as a convex mirror with focal length *r/*2, the virtual image lies at the midpoint of the chord connecting the LED and camera directions on the unit sphere:

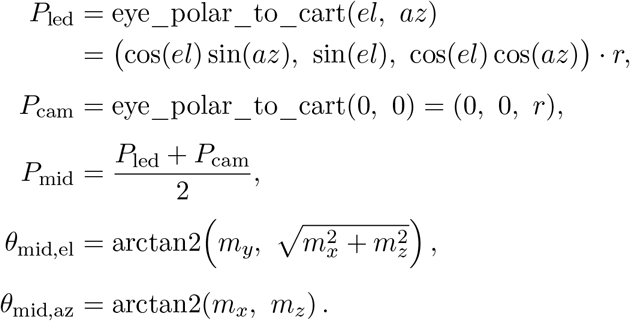

For small angles, this is well-approximated by *θ/*2, but the chord midpoint computation is exact for all angles. For the NE LED (*el* = −1.18°, *az* = −14.19°): midpoint_*el*_ *≈* −0.59°, midpoint_*az*_ *≈* −7.10°.

### A.6 Gaze angle computation from pixel coordinates

Given pixel coordinates of the pupil (*p*_*x*_, *p*_*y*_), LED reflection (*f*_*x*_, *f*_*y*_), and camera reflection (*c*_*x*_, *c*_*y*_), all displacements are measured relative to the camera reflection:

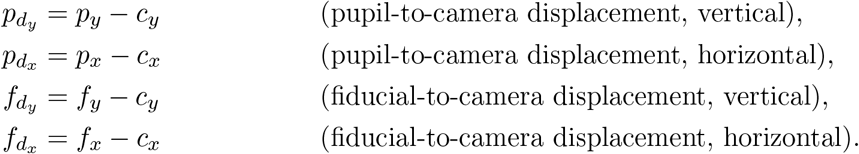

The camera reflection serves as the zero-gaze reference (optical axis). The ratio *p*_*d*_*/f*_*d*_ equals 0 when the pupil is at the camera reflection (gaze along the optical axis) and equals 1 when the pupil is at the LED reflection (gaze in the LED direction). The gaze angle in elevation is:

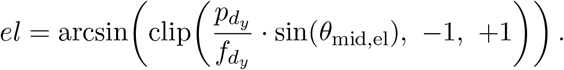

The gaze angle in azimuth (with foreshortening correction):

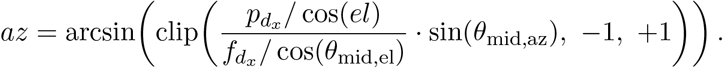

The cos(*el*) and cos(*θ*_mid,el_) terms correct for the foreshortening of horizontal displacements at non-zero elevation angles.

### A.7 Eye-to-table coordinate transform

The gaze direction in eye-centered spherical coordinates (*θ*_el,eye_, *θ*_az,eye_) is converted to table-centered coordinates as follows.

1. Convert eye spherical to eye Cartesian:

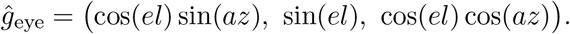
2. Rotate to world (table) Cartesian using the basis matrix:

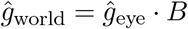

(since *B* transforms from world to eye, *B*^*T*^ = *B*^−1^ transforms from eye to world; here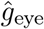 is a row vector).
3. Decompose into table-centered spherical:

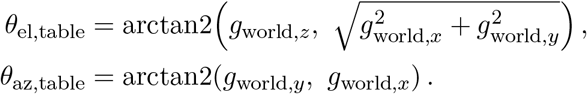

*Note:* the table convention uses *z* = vertical (elevation computed from the *z*-component), and azimuth is measured from the *x*-axis (lateral) toward the *y*-axis (depth/display) using the standard arctan2(*y, x*) convention.

