## Supplementary Methods, Supplementary Tables, Supplementary Figures for "Single-camera, calibration-free gaze estimation using corneal reflections"

#### S1. Hardware specifications

##### S1.1 Camera and optics

- Camera: FLIR Grasshopper3 USB3 NIR, part number GS3-U3-41C6NIR-C (Teledyne FLIR). 4.1 MP CMOSIS CMV4000-3E12 sensor, 1" global shutter, 90 fps maximum, C-mount. The stock IR cutoff filter is replaced with transparent glass.
- Lens assembly (Thorlabs, 0.7–4.5× magnification, 12 mm fine focus):
  - MVL6X12Z — 6.5× zoom lens
  - MVL20A — 2.00× extension tube
  - MVL6X025L — 0.25× magnifying attachment
  - MVLCMC — C-mount adapter
- Typical working distance: 5–15 cm from the mouse eye.

##### S1.2 Infrared illumination

- Fiducial LEDs: 850 nm, 5 mm through-hole, clear lens (Chanzon). Driven at ∼20 mA via 150–220 Ω current-limiting resistors.
- Flood lamps: Univivi 850 nm 4-LED arrays.

##### S1.3 Mounting

- Camera: articulating arm or post-and-clamp assembly. Each component's 3D position must be measurable to ±1 mm and remain fixed for the duration of a recording session.
- LEDs: 3D-printed holders or articulating arms; same requirements.

##### S1.4 Stimulus display and synchronization (electrophysiology validation only)

- Monitor: GTEK F2465P, 24", 1920 × 1080 IPS, 165 Hz.
- DAQ: LabJack T7 USB DAQ, 14 analog inputs, 16-bit, up to 100 kS/s.
- Neural recording: Neuropixels probes.
- Synchronization: photodiode mounted on the monitor, sampled by the DAQ alongside locomotion wheel and camera strobe signals.

##### S1.5 Computational requirements

- ≥8 CPU cores, ≥16 GB RAM, SSD storage (NVMe preferred). No GPU required.
- Python ≥3.9 with PyQt5, NumPy, SciPy, scikit-image, OpenCV, h5py, PyYAML.
- Tested on Ubuntu 24.04 LTS.
- Installation and current dependency list: `https://github.com/baccuslab/btrack/`.

#### S2. Geometry file format

Hardware positions are stored in a YAML configuration file. Each entry contains a label, a single-character tag identifying the component type, and the $(x, y, z)$ coordinates in centimeters in the world coordinate system. Recognized tags are: `E` (eye), `C` (camera lens), and `F` (fiducial LED). Example:


```
points:
  - {label: eye_headbar, tag: E, coords: [0.00, 0.00, 14.25]}
  - {label: cam_lens,    tag: C, coords: [-3.95, 3.97, 24.25]}
  - {label: led_ne,      tag: F, coords: [-2.07, 6.24, 24.25]}
  - {label: led_se,      tag: F, coords: [-1.40, 7.05, 11.75]}
  - {label: led_sw,      tag: F, coords: [-11.91, -5.67, 11.75]}
  - {label: led_nw,      tag: F, coords: [-9.40, 6.50, 24.25]}
```


Labels are free text; they do not affect the computation. Coordinates may be measured from any consistent origin, as they are recentered on the eye on loading. On loading, the software identifies the entry with tag `E` as the eye position (exactly one required), recenters all coordinates so the eye is at the origin, constructs the orthonormal eye basis from the eye-camera direction (Appendix A.2), and computes each fiducial LED's elevation and azimuth in the eye-centered spherical system (Appendix A.4). A geometry file with fewer than two fiducial LEDs is rejected.

#### S3. Image processing parameters

The pupil and fiducial corneal reflections are detected via a difference-of-Gaussians (DoG) bandpass pipeline applied independently to two channels: an inverted channel for the pupil (so the dark blob becomes bright), and a direct channel for the bright LED reflections. Connected components in the thresholded DoG output are filtered by area and selected by proximity to a user-defined reference point; the centroid of the selected component is taken as the sub-pixel position. Two blink criteria are available: a standard criterion flagging frames in which one or more fiducial reflections are displaced by more than 30 pixels (after 5-frame smoothing) with a pupil-area change exceeding 70% of the running median, and an alternate criterion using the area of the brightest infrared flood-lamp reflection. Detected blink intervals are post-processed by gap filling and linear extrapolation at boundary frames. When one fiducial LED is occluded but at least one other is visible, the occluded LED's pixel position is reconstructed using the inter-LED pixel offset measured from frames in which both LEDs are simultaneously detected.

#### S4. Manual curation

Following manual blink detection and trajectory processing, gaze traces can be manually reviewed to identify residual tracking artifacts, including missed blinks, false-positive blink classifications, and abrupt trajectory discontinuities. Suspect regions can be fixed in manual curation mode, which enables blink curation, interpolation across gaps, and extrapolation at blink edges through a graphical interface for frame-by-frame inspection and editing. Examples applications of these manual curation procedures are shown in Figure S4. An overview of the full three-panel GUI is shown in Figure S5.

### Supplementary Tables

**Table 1:** Hardware bill of materials (approximate 2025 USD prices).

| Component | Part number | Qty | Supplier | Price (USD) |
|---|---|---|---|---|
| *Camera & optics* | | | | |
| FLIR Grasshopper3 USB3 NIR camera (4.1 MP, 90 fps, 1” global shutter) | GS3-U3-41C6NIR-C | 1 | Teledyne FLIR | ~1,539 |
| 6.5× zoom lens with 12 mm fine focus | MVL6X12Z | 1 | Thorlabs | ~570 |
| 2.0× extension tube | MVL20A | 1 | Thorlabs | ~195 |
| 0.25× magnifying lens attachment | MVL6X025L | 1 | Thorlabs | ~130 |
| C-mount adapter | MVLCMC | 1 | Thorlabs | ~35 |
| | | | *Camera & optics subtotal* | ~2,469 |
| *Illumination* | | | | |
| IR LED, 850 nm, 5 mm through-hole (pack of 100) | — | 1 | Chanzon | ~10 |
| IR flood lamp, 850 nm, 4-LED array | — | 1 | Univivi | ~20–30 |
| LED resistors & breadboard | — | 1 | Digikey | ~5–10 |
| | | | *Illumination subtotal* | ~35–50 |
| *Mounting & positioning* | | | | |
| Articulating arm / post-clamp assembly | — | 1 | Thorlabs | ~50–150 |
| LED positioning arms / 3D-printed holders | — | 2–4 | Custom | ~10–30 |
| | | | *Mounting subtotal* | ~60–180 |
| *Data acquisition & sync (electrophysiology validation)* | | | | |
| LabJack T7 USB DAQ | T7 | 1 | LabJack | ~349 |
| USB 3.0 cable, Type-A to Micro-B (≥2 m) | — | 1 | Amazon | ~10–15 |
| | | | *DAQ & sync subtotal* | ~359–364 |
| *Stimulus display (electrophysiology validation)* | | | | |
| GTEK F2465P IPS monitor (24”, 165 Hz) | F2465P | 1 | GTEK | ~250–350 |
| | | | *Display subtotal* | ~250–350 |
| | | | **Eye tracker only (camera + optics + illumination + mounting)** | ~2,564–2,699 |
| | | | **Full system including DAQ + display** | ~3,173–3,413 |

**Table 2:** Default processing parameters for each processing stage.

| **Stage** | **Default parameters** |
|---|---|
| Pupil DoG | exp= 2, $\sigma_{\text{small}} = 20$, $\sigma_{\text{large}} = 50$, threshold= 210 |
| Fiducial DoG | exp= 1, $\sigma_{\text{small}} = 3$, $\sigma_{\text{large}} = 11$, threshold= 110 |
| Blink detection | LED threshold= 30, LED smoothing= 5, dPupil area= 0.7, blob area threshold= 70% |
| Artifact rejection | pupil jump= 15 px, min pupil area= 0.3, LED velocity= 5 px/frame, spike threshold= 2.0 deg/frame |

### Supplementary Figures

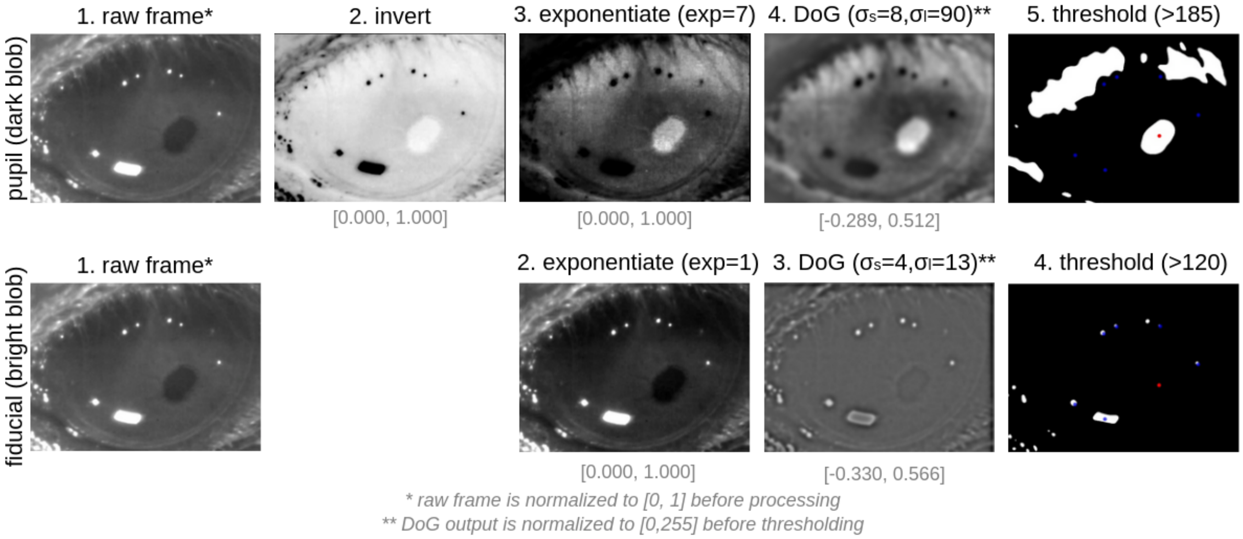


**Figure S1: Difference-of-Gaussians (DoG) detection pipeline.** (Top row) Step-by-step processing for the pupil channel: raw frame, inversion, exponentiation, DoG bandpass, thresholding, connected components, and final centroid selection. (Bottom row) Step-by-step processing for the fiducial channel (same pipeline without inversion).

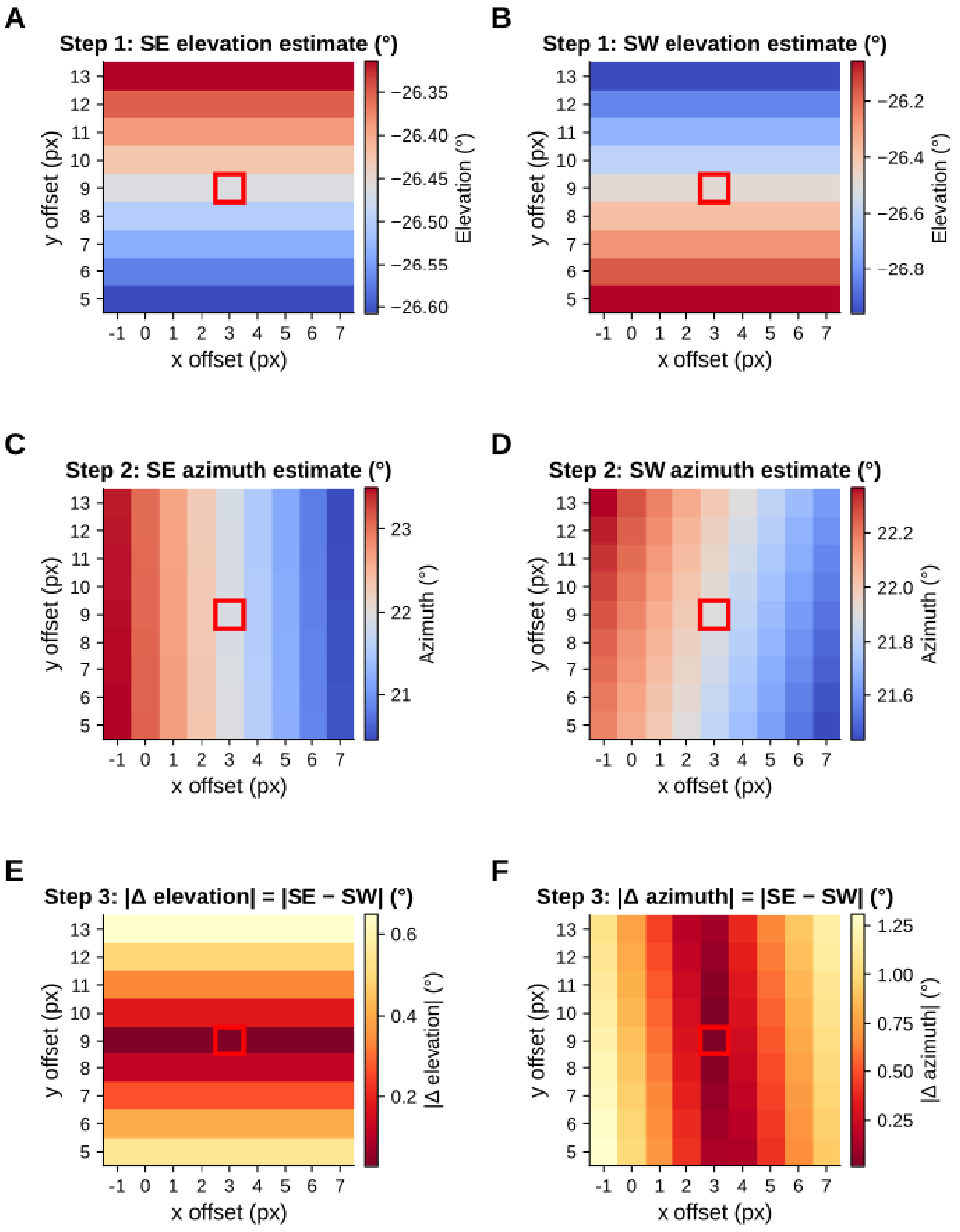


**Figure S2: Step-by-step grid search calibration.** Detailed view of a zoomed region around the optimal offset in the full $100 \times 100$ grid search. Each cell corresponds to a candidate pixel offset applied to the camera position estimate; the region shown spans $\pm 4$ pixels around the optimal solution in each axis (red box). (A–B) SE and SW elevation estimates at each candidate offset. (C–D) SE and SW azimuth estimates. (E–F) Absolute inter-LED disagreement: $|\Delta\text{elevation}| = |\text{SE} - \text{SW}|$ and $|\Delta\text{azimuth}| = |\text{SE} - \text{SW}|$. The optimal offset is selected by minimizing $|\Delta\text{elevation}|$ first, then $|\Delta\text{azimuth}|$ as a tiebreaker.

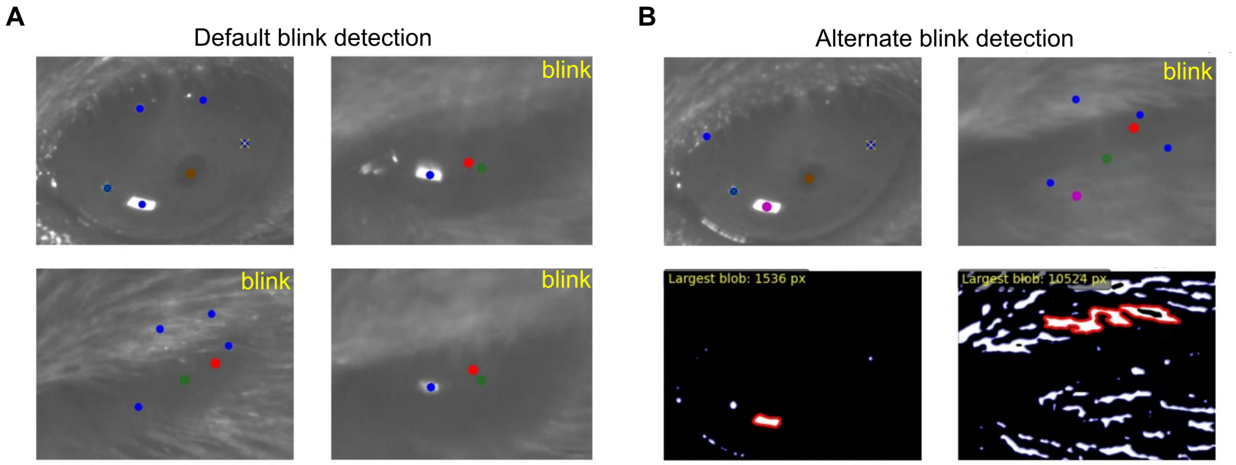


**Figure S3: Blink detection and fiducial correction.** (A) Standard blink detection: LED displacement trace (top) and pupil area trace (bottom) with detected blink intervals marked. (B) Alternate blink detection: reference LED blob area trace showing blink-associated area reductions below the threshold.

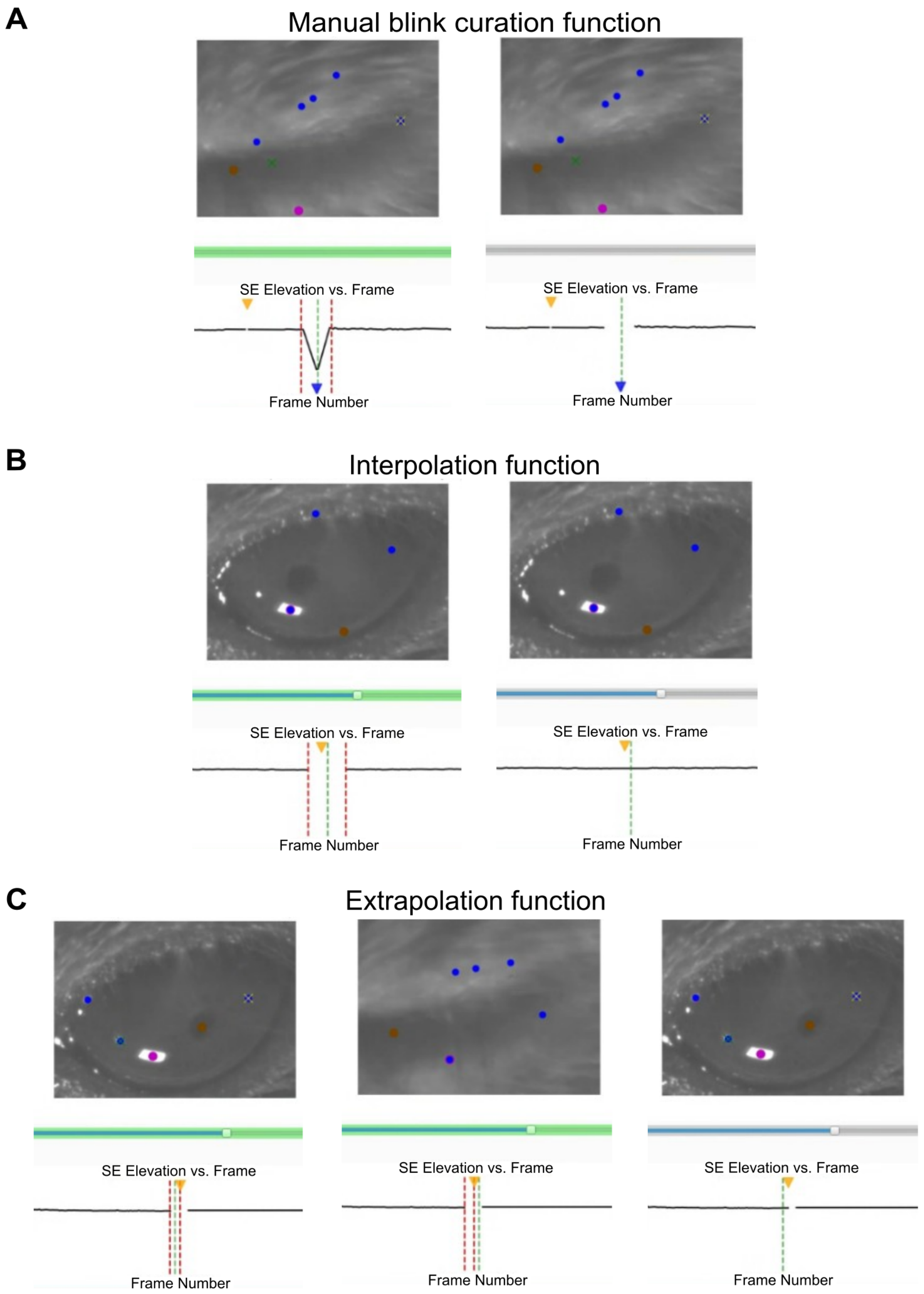


**Figure S4: Manual Curation Functions.** (A) Example manual blink curation function application to correct erroneous gaze traces with eye tracking inspection for gaze validation. Other example manual curation functions to correct miscellaneous erroneous gaze trace using (B) interpolation from both ends or (C) extrapolation from either end of a gaze trace window that requires correction.

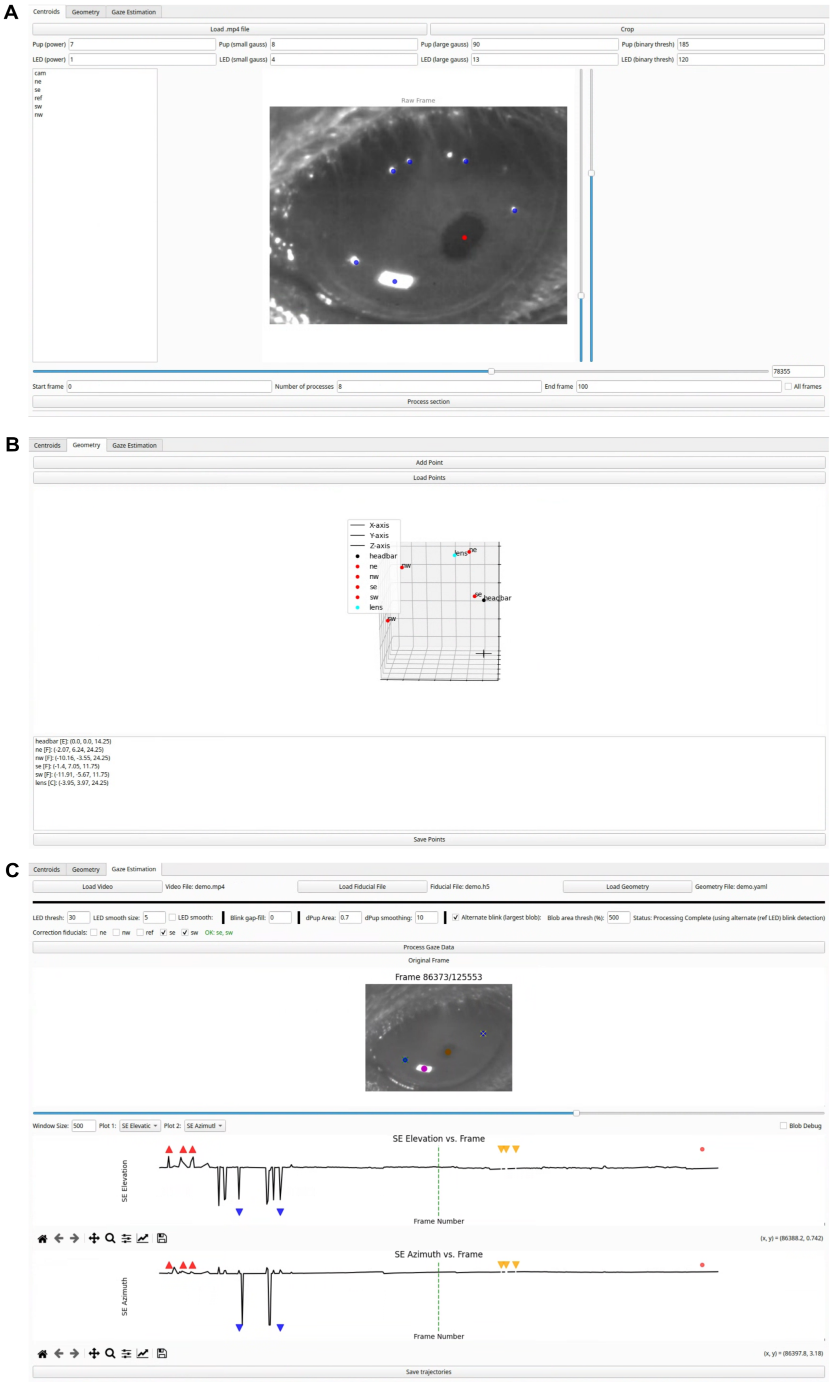


**Figure S5: Three-panel GUI Overview.** (A) Example view of the Centroids tab showing manually identified fiducial LED reflections (blue markers) and pupil position (red marker) used for pupil and LED detection via Difference-of-Gaussians filtering. (B) Geometry tab with an example experimental geometry file loaded, displaying the relative 3D geometry definition for eye center, fiducial LEDs, and camera positions used for gaze reconstruction. (C) Gaze Estimation tab for gaze angle computation displaying representative SE elevation and azimuth gaze traces with automatically detected anomaly markers overlaid, including spike events (red triangles), dip events (blue triangles), blink events (yellow triangles), and additional trajectory irregularities (red circles) used to guide blink detection, fiducial correction, and manual curation.
